# Evaluating FAIR-Compliance Through an Objective, Automated, Community-Governed Framework

**DOI:** 10.1101/418376

**Authors:** Mark D Wilkinson, Michel Dumontier, Susanna-Assunta Sansone, Luiz Olavo Bonino da Silva Santos, Mario Prieto, Peter McQuilton, Julian Gautier, Derek Murphy, Mercѐ Crosas, Erik Schultes

**Author notes:** Corresponding authors: Mark Wilkinson, Susanna-Assunta Sansone, Erik Schultes.

## Abstract

With the increased adoption of the FAIR Principles, a wide range of stakeholders, from scientists to publishers, funding agencies and policy makers, are seeking ways to transparently evaluate resource FAIRness. We describe the FAIR Evaluator, a software infrastructure to register and execute tests of compliance with the recently published FAIR Metrics. The Evaluator enables digital resources to be assessed objectively and transparently. We illustrate its application to three widely used generalist repositories - Dataverse, Dryad, and Zenodo - and report their feedback. Evaluations allow communities to select relevant Metric subsets to deliver FAIRness measurements in diverse and specialized applications. Evaluations are executed in a semi-automated manner through Web Forms filled-in by a user, or through a JSON-based API. A comparison of manual vs automated evaluation reveals that automated evaluations are generally stricter, resulting in lower, though more accurate, FAIRness scores. Finally, we highlight the need for enhanced infrastructure such as standards registries, like FAIRsharing, as well as additional community involvement in domain-specific data infrastructure creation.

## Introduction

The FAIR Data Principles[1] were designed to ensure that all digital resources can be Findable, Accessible, Interoperable, and Reusable by machines (and therefore, also by humans). The Principles act as a guide to the kinds of behaviours that researchers and data stewards should increasingly expect from digital resources[2], and in turn, the compliance-requirements on researchers when they publish scholarly outputs. However, despite their rapid uptake throughout the community, the question of how the FAIR principles should manifest in reality has been prone to diverse interpretation. As interest in, and support for the Principles has spread, this diversity of interpretation has broadened, in some cases to the point that some resources claim to “already be FAIR” - a claim that, currently, cannot be objectively and reproducibly evaluated. There was therefore a sense that the manifold interpretations and emerging implementations of FAIR pose a risk of fragmentation and siloing among and between digital resources in a manner antithetical to their intended purpose (or worse, being ignored entirely due to a lack of formal clarity).

Recently, the FAIRmetrics.org working group [http://fairmetrics.org] has published an open framework[3] for the creation and publication of metrics to objectively evaluate the many possible dimensions of FAIR that have been published. The framework was used to generate an initial set of 14 FAIR Metrics. Several communities, including the Global and Open (GO) FAIR initiative (GO-FAIR, http://go-fair.org), the US National Institutes of Health (NIH) Data Commons Pilot (https://commonfund.nih.gov/commons) are already testing and actively discussing to address the clarity, precision, and objectivity of these exemplar Metrics and the FAIR Principles to which they apply. The availability of FAIR metrics also necessitates a means of applying those metrics to evaluate the FAIRness of a variety of digital objects. We describe two approaches: i) a questionnaire for manual assessment, and ii) a semi-automated evaluation system.

Discussions with stakeholder communities highlighted some important behaviors or features that should be reflected in, or result from, the final output from this initiative. In particular:

1. Some metrics may be considered ‘universal’ but these may be complemented by additional resource-specific metrics that reflect the expectations of particular communities.
2. The metrics themselves, and any results stemming from their application, must be FAIR.
3. Open standards around the metrics should foster a vibrant ecosystem of FAIRness assessment tools.
4. Various approaches to FAIRness assessment should be enabled (e.g. self-assessment, task forces, crowd-sourcing, automated), however, the ability to scale FAIRness assessments to billions if not trillions of diverse digital objects is critical.
5. FAIRness assessments should be kept up to date, and all assessments should be versioned, have a time stamp and be publicly accessible.
6. FAIRness "level", presented as a simple visualization, will be a powerful modality to inform users and guide the work of producers of digital assets.
7. The assessment process, and the resulting FAIRness level, should be designed and disseminated to positively incentivise the digital resource, which should benefit from this and use it as an opportunity for improvement.

Here, we describe the FAIR Evaluator, an evaluation framework designed with the goal of addressing these community needs and requirements. The FAIR Evaluator is an open source, Web-based and machine-accessible framework that allows *de novo* publishing, discovery, assembly, and application of FAIR Metrics. FAIR Compliance Tests provide an objective, automatic, and reproducible output indicating the degree of compliance of the digital resource with a given Metric, and its associated FAIR Principle. The framework allows all stakeholders to evaluate the FAIRness of a resource, and importantly, allows data stewards to objectively and quantitatively monitor their progress when implementing their FAIR data stewardship plan. The Evaluator permits the *ad hoc* application of selected subsets of Metrics, the composition of which is optimized to serve the needs or interests of different communities. Evaluator outputs, which are FAIR themselves, enable discovery, comparison, and reuse of digital resources based on objective FAIRness evaluations.

The FAIR Evaluator framework is described in the Methods section. The Results section presents an overview of the result of applying the Metrics, both manually and via the Evaluator, to three resources, stored in Harvard Dataverse[4], Zenodo[5], and Dryad[6]. We have selected these generalist data repositories for the variety of their data content, as well as for their wider adoption by journals and publishers, as visible from their records in FAIRsharing[7] (Dataverse:https://doi.org/10.25504/FAIRsharing.t2e1ss; Zenodo: https://doi.org/10.25504/FAIRsharing.wy4egf; Dryad: https://doi.org/10.25504/FAIRsharing.wkggtx). We then discuss both the utility of the Evaluator, and some observations that were revealed by attempting to optimize the “FAIRness score” of a given resource, which are now being used to improve the precision and utility of the original Metrics set. Lastly, the automated FAIR Evaluator revealed gaps in our current data publishing practices that highlight the need for additional technological and organizational infrastructures to assist with FAIR compliance.

## Methods

The technical design considerations regarding an automated framework for a metrics-based FAIRness evaluation were:

1. Anyone should be able to evaluate anything.
2. Anyone can create and publish a metric or a collection of metrics.
3. An evaluation is performed for a particular resource using one or more metrics.
4. All components of the evaluation framework should themselves be FAIR, including the result of the evaluation.
5. The evaluation should be as objective as possible (i.e. the code should avoid accommodating “special cases”).
6. The evaluation should include clear feedback for improvement.
7. The evaluation framework is modular and reusable.
8. The framework should automatically update to accommodate new standards or technologies whenever possible, to minimize maintenance and sustain utility over time.

Thus we pursued a design framework that allowed the public definition of, and collection of, Metrics that were relevant to their community, and an interface that would allow that collection to be applied to any given resource.

### Overall design

The evaluation framework (the “Evaluator”) is a Ruby on Rails application with three primary components:

- A (read/write) registry of known Compliance Tests (defined below)
- A (read/write) registry of collections of Compliance Tests
- A registry of evaluations, including the resource evaluated, input provided, and output results

### Definition and registration of Compliance Tests

Fourteen FAIR Metrics were defined[3]. A template to create new metrics is also included in that work in order to help communities develop additional metrics. Thus, our framework is open to the registration of novel, community-designed metrics, so long as that community simultaneously creates an associated Compliance Test (Figure 3, left panel)

A Compliance Test is a Web-based software application that tests compliance of a Resource against a given Metric. Compliance Test interfaces must be described using a smartAPI (Swagger/OpenAPI-based) interface definition, with rich FAIR metadata describing the interface. The URL to this smartAPI[8,12] definition is presented to the Evaluator, which then automatically extracts the metadata about the interface, and the FAIR Principle being tested, in order to complete the registration of this new Compliance Test. There is not a 1:1 mapping between Metrics and Compliance Tests; some Compliance Tests might only examine one facet of a Metric, thus requiring additional Compliance Tests to achieve a full representation of the Metric. Also, certain Metrics (for example, FM_R1.3) were considered difficult to test, and thus no Compliance Test has been coded to-date.

### Definition and registration of Metrics Collections

Different communities may have dramatically different interests with respect to assessing the FAIRness of a resource; for example, a funding agency may want to ensure that the metadata acknowledges its participation, while a data integration project may be primarily interested in data-level interoperability. As such, The Evaluator allows the assembly of individual Compliance Tests into a Metrics Collection (Figure 1, right panel). These collections are tagged with metadata related to the community for which they might be considered relevant (however, this does not preclude their use by other communities).

**Figure 1:**
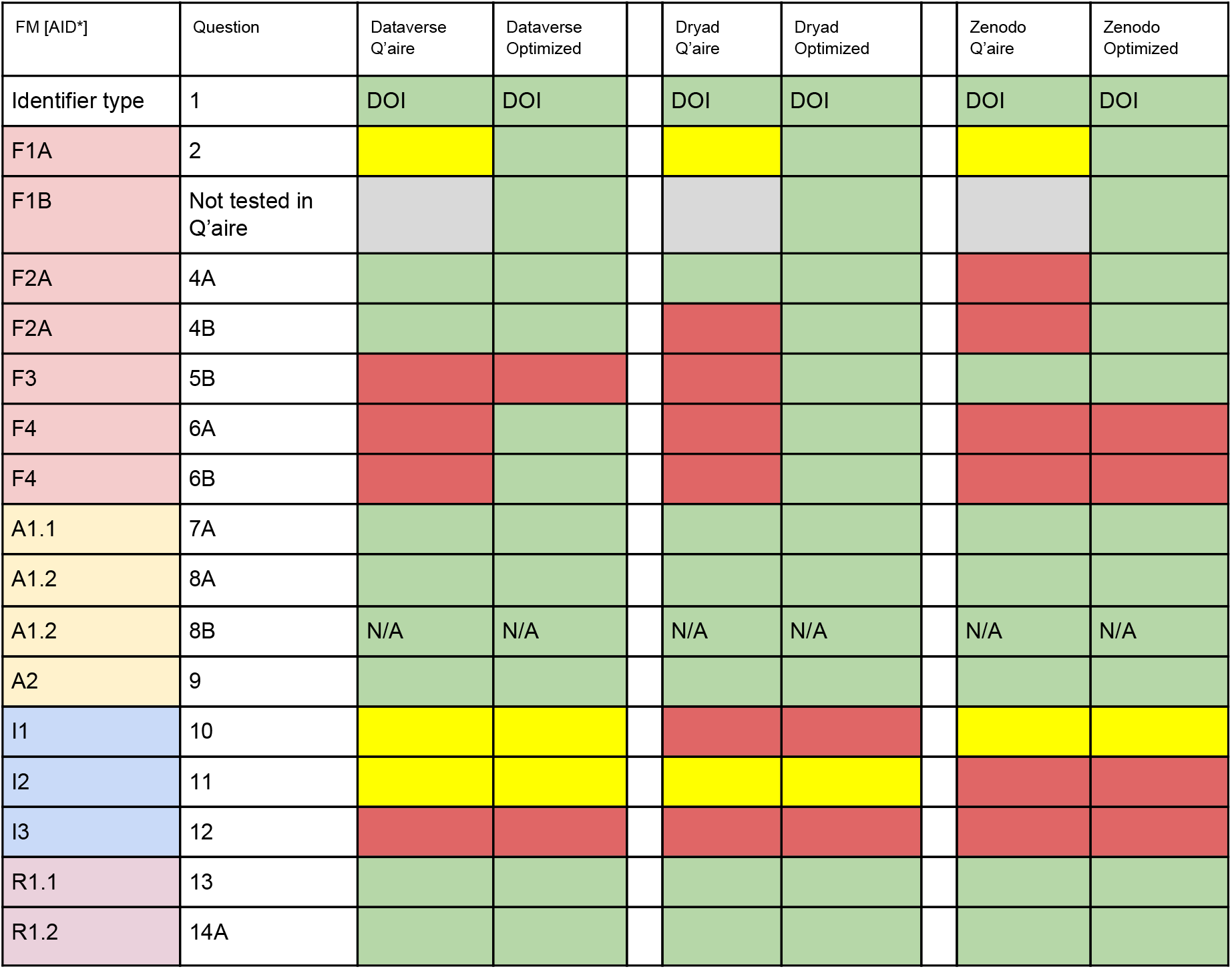
Evaluator (automated) scoring of original, non-optimized questionnaire responses compared to optimized responses provided by FAIR Data experts. Green indicates that the metric was considered a “pass” by the human evaluators, or the Evaluator software. Yellow indicates an intermediate level of success (less than perfect score) by the Evaluator software, or cases where the human evaluator considered the evaluation to be flawed or where the input required manipulation in some way (see detailed annotations in the supplementary material). Red indicates a score of zero/fail by both humans and/or the Evaluator software. DOI is the “canonical-form” DOI (not a Qname, not a URL) for that resource. Metric R1.3 was not scored, as we consider its expectations impossible to meet at this time. The responses to questionnaire F1B were not scored, as we determined that the question was misleading.

Collections of metrics are published with their own metadata as in the form of a ‘FAIR Accessor’[13] - a metadata document modelled on the W3C’s Linked Data Platform specification for ‘collections’[11]. The FAIR Accessor, therefore, contains the metadata about that specific Metric Collection, in addition to pointers to the individual Metrics contained (via their smartAPI URLs)

### Application of a Metric Collection to a Resource

Testing the compliance of a resource against a Metric Collection is called an “Execution”. In the Evaluator, when a Metric Collection is displayed, there is a button to begin a new Execution. Activating an Execution (Figure 4) causes the Evaluator to retrieve all smartAPI descriptions from all of the Metric Testers associated with that Collection. It parses these and creates a single Web Form representing the combination of these interfaces, together with a field that represents the GUID of the Resource to be evaluated.

Each FORM field is labelled with the type of data required by that smartAPI interface - for example “what is the URI of the registered identifier schema used by this Resource?”. Currently, these fields are filled-in manually. Once all fields are complete, the FORM data is passed to the Evaluator, which separates each FORM value into a correctly-formatted input to its associated Metric Tester smartAPI. The Evaluator collects the output of each test (so far, only boolean or percentage scores), and collates these into a unified output, which in this iteration uses a “five star” system. Each Execution has recorded metadata, including the inputs that were sent, and the output returned. This will allow, for example, a data host to monitor their degree of improvement over time, as they attempt to become more FAIR, by simply re-executing the Evaluation. It also allows public access to the output of the evaluation, such that individuals or agents could make decisions regarding, for example, whether or not they wish to use a given data resource, based on its FAIRness evaluation.

### Additional Resources Required

To successfully create the Evaluator, the requirement for additional FAIR infrastructure became apparent. In both our manual and automated evaluations, we noted that many of the questions we asked had no ‘canonical’ answer by which we could evaluate the validity of the response. For example, what is the ‘canonical’ URI that represents the DOI identifier schema? This issue arose repeatedly with respect to various metrics such as identifier schemas and their persistence, machine-readable data formats and file formats, knowledge-representation languages, licenses, policy documents, and so on. In these cases, it is currently not possible to validate the information provided by the user of the Evaluator, beyond simply confirming that the URL they provide does, in fact, resolve.

We are working with FAIRSharing[7] to register additional elements such as transport protocols, generic file formats and other community standards required by the FAIR Metrics Tests but that currently lack ‘canonical’ URIs to represent them. The work is ongoing, also as part of FAIRsharing’s role in several international infrastructure programmes; additional (or more robust) Compliance Tests will become available in-parallel with the creation of these new FAIRSharing resources. In the interim, several of the existing Compliance Tests will always respond with a “pass”, regardless of input, making the current evaluation scores somewhat artificial; however, we want to bring this resource to the attention of the community now, regardless, as we are requesting broad discussion and input into the creation of this important piece of FAIR Data infrastructure.

## Results

The Metrics used in this study are briefly described in Table 1. Our results are summarized in Figures 1 and 2. The full overview of results, annotations related to those results, and links to the detailed inputs and outputs to/from the Evaluator, are available as online supplementary data (Data Citation 1). There is also an ongoing public discussion regarding the clarity and utility of the Metrics, and the methodology for testing compliance using the “Issues” functionality of GitHub (https://github.com/FAIRMetrics/Metrics). A machine-readable metadata document, including links to individual Compliance Tests, is available at http://linkeddata.systems/cgi-bin/fair_metrics, with a human-readable equivalent available through the FAIR Evaluator interface, through the smartAPI Registry (search term “FAIR Metrics”) or by opening the Compliance Test metadata in the smartAPI Editor linked from the smartAPI Registry (e.g., http://smart-api.info/editor/f6a4c1321e0b86746ad229e06713a635).

**Table 1:** The current set of FAIR Metrics[3].

|  |  |
| --- | --- |
| FM_F1A | Whether there is a scheme to uniquely identify the digital resource. |
| FM_F1B | Whether there is a policy that describes what the provider will do in the event an identifier scheme becomes deprecated. |
| FM_F2 | The availability of machine-readable metadata that describes a digital resource. |
| FM_F3 | Whether the metadata document contains the globally unique and persistent identifier for the digital resource. |
| FM_F4 | The degree to which the digital resource can be found using web-based search engines. |
| FM_A1.1 | The nature and use limitations of the access protocol. |
| FM_A1.2 | Specification of a protocol to access restricted content. |
| FM_A2 | The existence of metadata even in the absence/removal of data |
| FM_I1 | Use of a formal, accessible, shared, and broadly applicable language for knowledge representation. |
| FM_I2 | The metadata values and qualified relations should themselves be FAIR |
| FM_I3 | Relationships within (meta)data, and between local and third-party data, have explicit and 'useful' semantic meaning |
| FM_R1.1 | The existence of a license document, for BOTH (independently) the data and its associated metadata, and the ability to retrieve those documents |
| FM_R1.2 | That there is provenance information associated with the data, |
| FM_R1.3 | Certification, from a recognized body, of the resource meeting community standards. |

As we noted in our earlier publication regarding the Metrics [3], measuring FAIRness, even using an automated system, remains to some degree “trust-based”; for example, the *quality* of a persistence policy narrative cannot be evaluated, rather, only the existence of the document can be verified. Nevertheless, to demonstrate the utility of the FAIR Evaluator, data stewards were provided a questionnaire covering all of the Metrics, and asked to provide responses pertaining to the resource of their choice within the repository system they represented. Their answers were (informally) evaluated[9] by two members of the FAIR Metrics Group. The evaluation was intended both to provide insight into the quality of the metric itself (i.e., was the question clear and answerable?), as well as to examine how the participating data stewards perceive what constitutes compliance with a Metric, compared to the perception of those with more extensive experience in FAIR infrastructures. The evaluation team then undertook to improve the responses, through a close manual examination of the data infrastructure being evaluated. For example, the documentation of the resource was examined to identify possible alternative data entry-points that might be more successfully interrogated by a machine; the HTML source of the Web page was examined by-eye to observe the type/structure of the identifiers being used in the HTML code, and if there were embedded metadata not visible in the browser; HTTP Response headers were examined to determine if alternative representations of the data were available; URLs were suffixed with a variety of common extensions (e.g. .rdf, .xml, ?xml, ?format=xml), since these are often used by data providers as a means of retrieving structured data *in lieu* of HTML; and URIs were interrogated using a variety of HTTP Accept headers to determine if the server would respond with more structured or machine-readable data. Based on this deep exploration of the resource, we selected the Evaluator inputs that appeared most likely to pass the majority of tests and re-evaluated the resource (full results are available in the supplementary material). In addition, although every resource in the questionnaire evaluation provided a URL as the answer to question 1 (“What is the IRI of the resource being evaluated?”), we changed this to the equivalent DOI for that resource. The primary reason for this is that, for the three resources selected for further evaluation, using a URL as the seed identifier resulted in numerous additional metrics failing, which reduced the utility of this study. Additional details related to this decision are presented in the Discussion section.

In Figure 1, we show a comparison of the Evaluator being applied to the original questionnaire responses from the data stewards, and to the “optimized” responses selected by the FAIR metrics authors after their deep inspection of the data resource. The most notable observation that can be drawn from this is that the same resource may show markedly different (i.e. better) FAIRness results based on what inputs are selected for evaluation (e.g. which GUID is used to initiate the evaluation) - an important consideration for data stewards who are attempting to optimize their evaluation scores.

To examine if and how automated evaluations differ from manual evaluations, we applied the Evaluator to the non-optimized questionnaire responses, with the caveat that in every case, we used the DOI of the resource being evaluated as input to the Evaluator, rather than the URL of that resource provided to us by the questionnaire responders. We then compared those automated evaluation results against the manual evaluation executed by the FAIR Metrics authors. The summary in Figure 2 shows that, generally and as expected, automated evaluation is more strict than manual evaluation. The few cases where the automated test passed, yet the manual test failed, generally represent cases where the evaluation is based on “trust”. For example, the automated test for a metadata longevity policy only attempts to resolve the URL it is given; it does not (at this time) attempt to validate the content of the document returned. It is in such cases where the human evaluator may indicate failure regarding that metric, whereas a machine cannot make such a qualitative determination.

**Figure 2.**
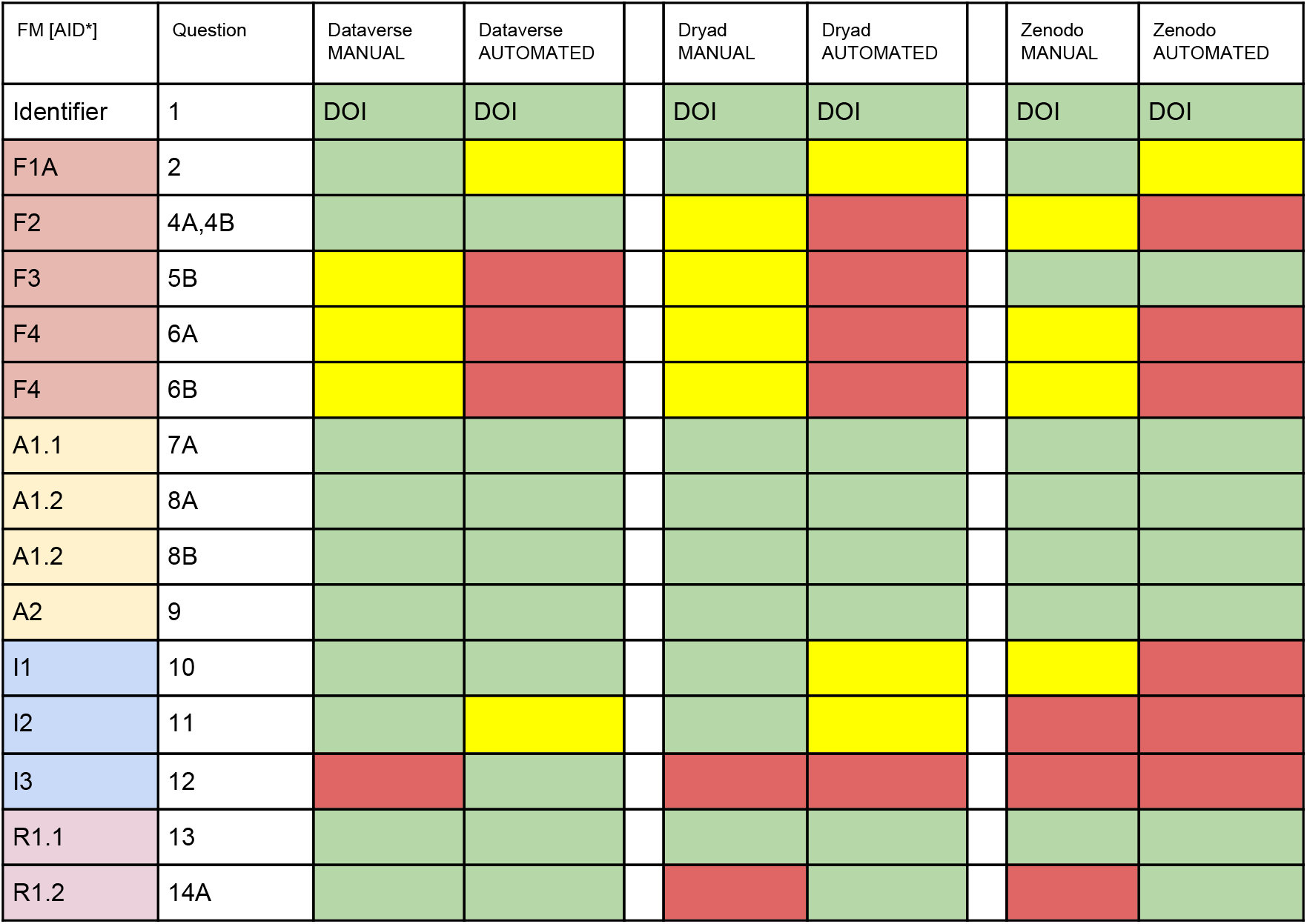
Comparison of Evaluator (automated) vs. manual scoring of non-optimized questionnaire responses. Green indicates that the metric was considered a “pass” by the human evaluators, or the Evaluator software. Yellow indicates either concerns expressed by the human evaluators, or an intermediate level of success (less than perfect score) by the Evaluator software. Red indicates a “fail” as assessed by the human evaluators, or a score of zero from the Evaluator software. DOI is the “canonical-form” DOI (not a Qname, not a URL) for that resource. Note that the the automated evaluation is incomplete compared to the questionnaire evaluation, because there are several Metrics that do not yet have a Compliance Test.

**Figure 3:**
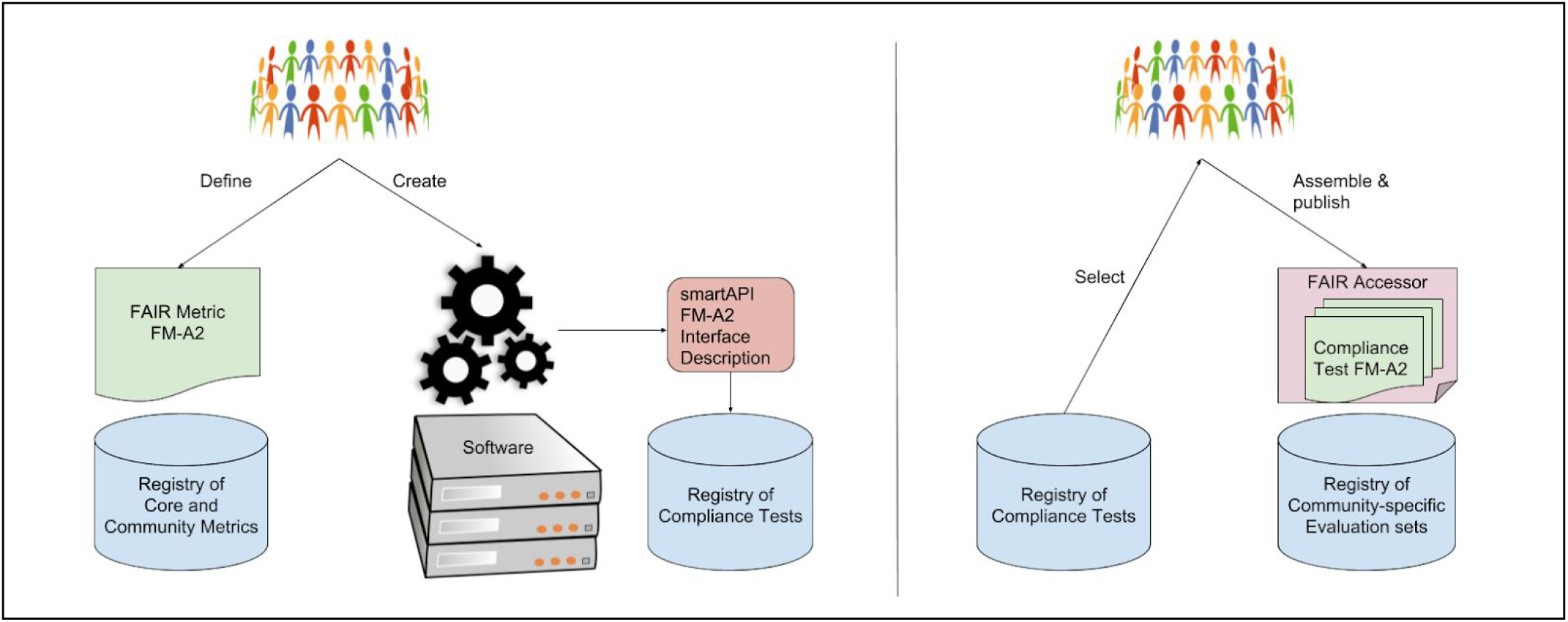
The community-driven process of creating and assembling FAIR Metrics, Compliance Tests, and Assessments. In the left panel, the community uses the Metrics Template to define a new FAIR Metric (for example, a metric to test FAIR Principle A2). This is registered in the registry of Metrics together with its metadata. They then create a piece of Web-accessible software capable of testing that metric - a Compliance Test. They describe the interface of that software in a FAIR manner (using smartAPI metadata), and the Compliance Test metadata is also published in a registry. In the right panel, communities select Compliance Tests for Metrics of interest from the registry, reflecting the expectations of FAIRness for their community These are assembled into a FAIR Accessor container representing an Evaluation metrics set, together with descriptive metadata about, for example, the applicable community for this metrics set, the authorship, etc. This is then deposited in a registry such that the Evaluation can be re-executed over any number of applicable resources.

**Figure 4:**
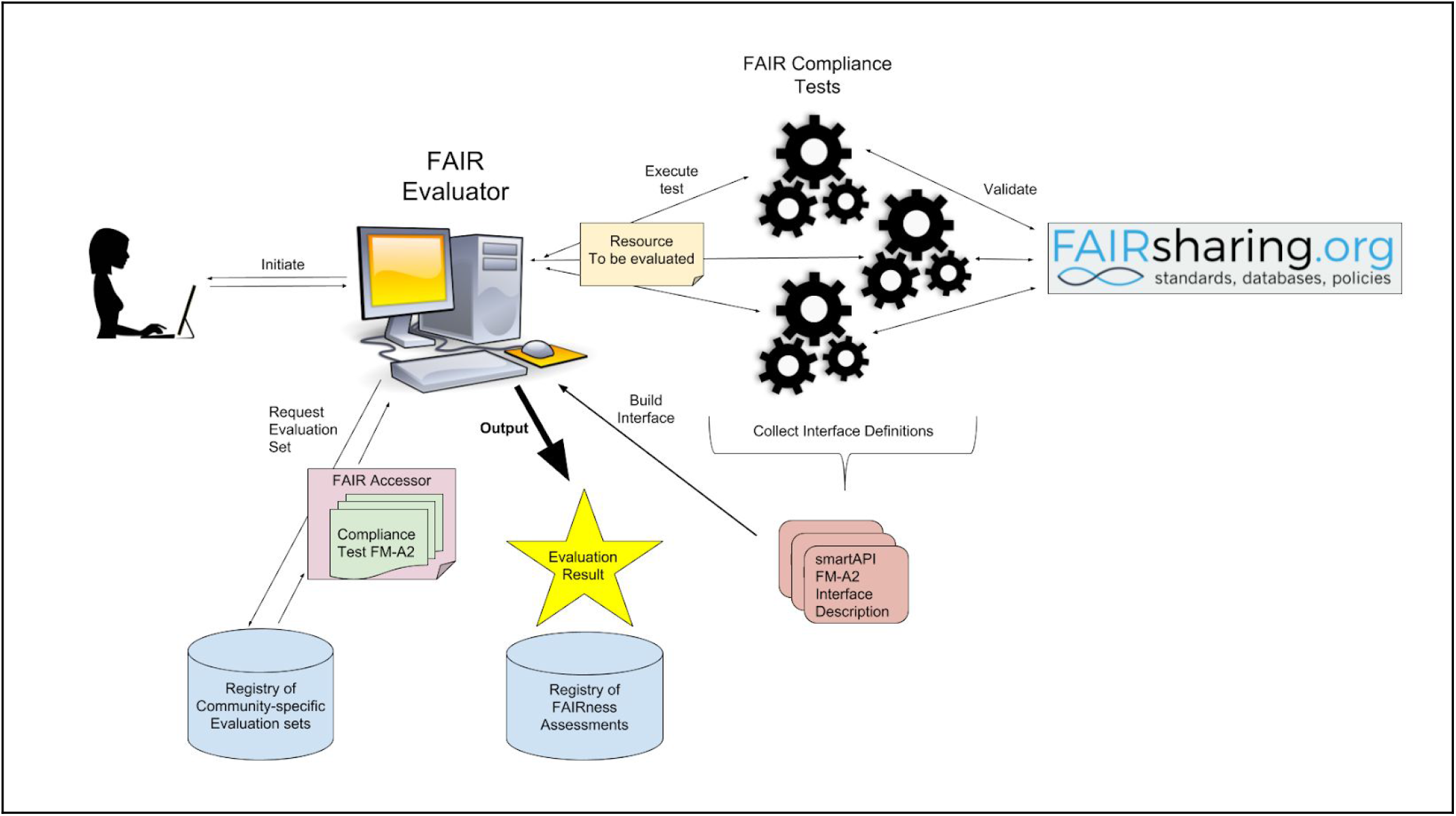
Using the FAIR Evaluator to evaluate a Resource. A user engages with the FAIR Evaluator to query the Registry of Evaluation sets to discover an appropriate set of Compliance Tests for the type of Resource that will be evaluated. Evaluation Sets contain metadata for each constituent Compliance Test, including the URL of the Compliance Test software’s smartAPI interface definition. These are retrieved via HTTP GET, returning the smartAPI descriptor of each Compliance Test. These component Compliance Test interfaces are are combined into a single HTTP FORM, or JSON API, by the Evaluator. The Resource to be tested, along with other required inputs to this test set, are passed to the relevant individual Compliance Tests, that in turn may validate the input data against standards registered in FAIRsharing. The outputs from all Compliance Tests are once again re-assembled into a single Evaluation Result, and this is stored in a registry of FAIRness Assessments. These FAIRness assessment scores can then be used by, for example, Web widgets, to quickly display the FAIRness level for any given Resource, in order to assist in the selection of highly-FAIR Resources, or to monitor progress towards FAIRness within a Data Management Plan.

## Usage Notes

Code for the FAIR Evaluator is available in GitHub (https://github.com/FAIRMetrics/Metrics/tree/master/MetricsEvaluatorCode). A Python client for the FAIR Evaluator is available in an independent repository (https://github.com/wilkinsonlab/FAIR_Evaluator_Client/). The public interface to the prototype Evaluator software is available for anyone wishing to execute evaluations (https://github.com/FAIRMetrics/Metrics/tree/master/MetricsEvaluatorCode) though certain functionality has been disabled to ensure stability during peer review.

## Discussion

With an increasing number of agencies and journals requiring FAIRness from their participant communities, and with claims to existing FAIRness being made by those participants, there was an urgent need to find a flexible, transparent, and objective way to measure the degree to which a given resource adheres to the FAIR Principles. The evaluation framework described here is one solution to that problem.

Flexibility is achieved by allowing community-participation in the design of Metrics, and Compliance Tests, through having an open design rubric and interface registry, respectively. Thus, if a particular aspect of FAIRness is not being measured by the existing Metrics set, a new Metric may be designed following the rubric, and a formal test of that Metric may be published on the Web and registered with our evaluation framework. Flexibility is also achieved in allowing different stakeholder communities to select the Metrics that are most relevant to them - ensuring that no resource is ‘unfairly’ tested against a requirement considered irrelevant by that community.

Transparency is achieved by making concrete Compliance Tests, using challenges that have clear ‘pass/fail’ responses. Moreover, examples of success and failure are provided as part of the (human-readable) Metric to further aid in understanding the purpose of the Metric, precisely what will be evaluated, and how.

Objectivity is achieved by mechanizing the evaluation of the stakeholder’s answers - ask, but verify. This is a notable aspect of the system for an additional reason; that while FAIRness is intended to apply to both humans and machines, it is the latter that is of greatest importance as we deal with an increasing volume and distribution of data and tools. We increasingly expect our computational agents to discover, integrate, and interpret data on our behalf. As such, this tool measures (primarily) FAIRness for machines, which is also the most challenging aspect of being FAIR. Moreover, when data is FAIR for machines, there are myriad tools that can assist in automatically making it FAIR for humans.

It is important to emphasize that FAIRness is a measure reflecting a continuum of data resource attributes and behaviours that result in increasingly re-usable data objects - one cannot “be FAIR”, only more or less FAIR. With this in mind, we note that these initial 14 Compliance Tests, and their associated Metrics, do no more than scratch the surface of evaluating the objectives envisioned by the FAIR Principles; a perfect score on these tests will only indicate that a commendable level of FAIRness has been achieved, not that a resource “is FAIR”. We fully anticipate that, over time, more challenging Metrics and Compliance Tests will be designed that, for example, eliminate the need for manual question-answering, since this is antithetical to one of the FAIR Principles’ original motivations that a machine should be able to discover and use data unaided.

Both the manual and automated analyses undertaken here highlighted gaps in existing data publication infrastructure - in particular, the need for additional standards, and a registry of those standards. For example, Compliance Test F1 requests a link to the specification of the formal identifier schema being used by the resource, and must also determine whether that identifier system guarantees global uniqueness. However, it isn’t clear what the “correct” link is in the case of, for example, a DOI. Should it link to the DOI project homepage, the PDF document that describes the DOI structure, or the Wikipedia entry describing DOIs? Even if this were agreed-upon, how would a machine determine if that specification guaranteed global uniqueness? In response to this emergent need, the FAIRSharing registry is beginning to create indexes of, and provide globally unique identifiers for, a wide variety of standards and policies that are relevant to FAIR evaluations. The first of these - an index of identifier schemas available to create Globally Unique Identifier (GUIDs) for data - became available during this analysis. These records in the FAIRsharing registry are now being used as the basis for automatically evaluating FAIR Principle F1 (globally unique and resolvable identifiers). Use of the FAIRsharing registry not only harmonizes and centralizes the representation of the diversity of global standards (identifier schemas, as well as reporting requirements, models/formats, terminology artifacts to represent and share data), but it also allows the Compliance Test to automatically evolve in parallel with the appearance of new standards, so that the Compliance Test itself does not deprecate - one of the key desiderata of our automated evaluation system.

Several additional observations are particularly noteworthy, since they were encountered in almost every case and thus serve to provide insight into ‘systemic errors’ regarding how we build our interfaces. They also guide us in our considerations when building data architectures aimed at mechanized exploration.

First, in every evaluation, the optimal identifier for a resource was its ‘canonical-form’ DOI - that is, the non-URL form, and without the ‘doi:’ prefix. From the perspective of automated exploration, the ‘canonical-form’ DOI matched all other DOI structures, and therefore allowed software to locate the identifier within a variety of different data serializations. The use of a URL as a the primary identifier often caused failures in identifier discovery due to the utilization of relative URLs, rather than absolute URLs, within HTML-based metadata documents. While, clearly, the process of generating absolute URLs from relative URLs in HTML is well-defined, this is primarily applied to browsing. The FAIR requirement (Principle F3) that the data identifier be explicitly present in the metadata is for the purpose of search, not navigation, and search engines do not (necessarily) fill-in relative URLs when indexing an HTML document. Thus, the use of relative URLs initially led to failure of the “data identifier in metadata” (Metric FM_F3) Compliance Test when a URL was proposed as the primary identifier for the record. After further consideration, the Compliance Test has now been updated to both expand all relative URLs prior to searching for the identifier, and to search for URL-encoded representations of the identifier in the HTML document; however, it may be argued that one or both of these is contrary to design consideration #5 (avoid accommodating “special cases” - See Methods section). An equally problematic observation related to the inconsistent use of the primary identifier throughout the interface. Even when a DOI was the primary identifier, there were often other identifiers used in various components of the interface, particularly in pages resulting from keyword searches where local/repository-specific URLs representing the data - not containing the DOI at all - were much more likely to be used to represent the discovered record. This led to failures in the Compliance Tests related to discoverability of a resource via keyword search.

Second, in every case evaluated, the DOI resolved to a URL that led to a Web page (i.e. HTML). These URLs did not, in our tests, respond to content-negotiation, though structured data was often provided through bespoke “download as” links that could not be automatically discovered. In all cases the default Web pages contained embedded structured metadata as JSON-LD. The metadata from Zenodo was relatively complete compared to the human-readable HTML, and included links to the data via the Schema.org DataDownload property; however, these downloads are not distinguished from one another by any additional metadata regarding the individual file content. In the case of Dryad and Dataverse, the embedded metadata was incomplete, consisting primarily of elements such as keywords, citation information, and narrative abstract metadata, but lacking explicit download links or references to the data being described, thus being non-compliant with Principle F3 (data identifier is explicitly part of the metadata). The paucity of structured (meta)data after resolution of the resource’s primary GUID is troubling, and highlights the continuing focus on infrastructures created for humans versus those created for machines.

Third, all three questionnaire respondents suggested Google as the search engine that would lead to discovery of their resource. While this was true, Google’s Terms of Service indicate that “[Google] do[es] not allow the sending of automated queries of any sort to our system without express permission in advance from Google.” We found no update to these terms as a result of the recent release of the Google Dataset Search tool (https://www.blog.google/products/search/making-it-easier-discover-datasets/). As such, Data Stewards may not directly rely on Google to achieve the discoverability aspects of FAIRness - where the resource must be discoverable by machines, as well as humans. In the three cases tested, the resource providers all had their own bespoke keyword search capability, and thus we were able to optimize their scores by using their own search functionality, rather than Google, to pass that Compliance Test.

A number of barriers remain, and many of these - for example, the design and adoption of new standards, and verification of community best-practices - cannot be overcome by individual data publishers; rather, they require coordination and effort within stakeholder communities. We hope that the desire to maximize the FAIRness level of a resource will spur various stakeholder communities to take-up these challenges and create the necessary standards, practices, and infrastructures.

It is worth reiterating that this study is primarily aimed at evaluating the current set of FAIR Metrics, and their instantiation as Compliance Tests; the evaluations executed here are not, in any way, an attempt to judge the repositories used in the study. Moreover, we point-out that several Compliance Tests evaluate aspects of a data deposit that are clearly outside the domain of responsibility of the repository host - for example, the provision of a formal linkset describing the internal and external linkages present in the data submission. Nevertheless, data publishing is a ‘collaboration’ between the depositor and the repository they select. We are aware that these kinds of evaluations will affect how the community perceives and/or judges host repositories, and thus it is useful to engage with these key stakeholders in a public discussion about the pros and cons of assigning “scores” to data resources, the appropriateness of individual metrics, and the “fairness” of the automated Compliance Tests. We requested feedback from leaders of the three high-profile repositories described in this study to learn their perspectives on the utility, rationality, and validity of this approach to assessing FAIRness, and how such evaluations might affect their operations in the future.

There was a certain degree of surprise expressed by these participants with respect to what a FAIRness test measured, and how. We believe this stems from the implementation ambiguity of the FAIR Principles, which aim to be a set of guideposts spanning an extremely broad range of data publishing scenarios. This was, in fact, a key objective of this work - that is, to make the Principles much more concrete and practical, and clarify expectations among data stewards of all types. In this sense, all participants felt that the exercise was useful.

A second common observation related to the Metric Test’s strong requirement for purely mechanized exploration. While this has always been the primary objective of FAIR, as indicated in the original FAIR Principles publication[1], and subsequent clarification[2], data stewards will generally lack the means for testing their degree of FAIRness for machines because, although structured metadata harvesting has achieved a high level of efficiency and utility (e.g. DataCite[10]), there is very little (publicly available) technical infrastructure aimed at fully mechanized, precise, and fine-grained data exploration, discovery, and integration in response to specific needs. Thus, FAIR Compliance Tests aid data stewards by giving them a transparent, objective tool to evaluate those facets of FAIRness we anticipate will be required to support fully mechanized exploration and integration. Moreover, the rigor of these tests will increase over time, as the published test requirements become more sophisticated, providing data stewards a stepwise evolutionary path towards increasingly “machine-actionable” data publications.

Finally, there was support for the decision to open-up the creation of Metrics, Compliance Tests, and Evaluation Sets to community input and definition. We considered this a core requirement given the diversity of data resources that should be FAIR, the diversity of communities that rely on those resources, and the diversity of stakeholders - including funding agencies, librarians, scholarly publishers, and data scientists - who are interested in different aspects of FAIRness.

FAIR is aspirational. The availability of a tool that objectively measures FAIRness allows both data owners, and third-party stakeholders, to track progress as the global community continues to move all digital resources towards an increasing degree of interoperability and reusability.

The FAIR Compliance Tests used in this analysis are continuously evolving, and are therefore made publicly available via a dynamically-generated machine-readable Linked Data Platform [11]-compliant container at http://linkeddata.systems/cgi-bin/fair_metrics. A ‘sandbox’ version of the Evaluator, with minimal layout or formatting, is available for public evaluation by following links provided at (https://github.com/FAIRMetrics/Metrics/tree/master/MetricsEvaluatorCode). This interface provides both human and machine interfaces, via a Web page, or a JSON-oriented REST API, respectively, and a Python library is available to interact with this API. Several functions have been hidden for the purposes of maintaining stability during review; full functionality, including the ability to execute new evaluations, is available on request to the authors. We specifically invite the community to participate in discussions about the Metrics and their implementation as Compliance Tests (https://groups.google.com/forum/#!forum/fairmetrics) to ensure that they reflect the perspectives of a wide range of stakeholders.

## Acknowledgements

MDW is funded by the Isaac Peral/Marie Curie cofund with the Universidad Politécnica de Madrid, and Ministerio de Economía y Competitividad grant number TIN2014-55993-RM. S.A.-S., P.MQ. and elements of FAIRsharing are funded by grants awarded to S.A.-S. from the UK BBSRC and Research Councils (BB/L024101/1; BB/L005069/1), EU (H2020-EU.3.1, 634107; H2020-EU.1.4.1.3, 654241; H2020-EU.1.4.1.1, 676559), IMI (116060), NIH (U54 AI117925; 1U24AI117966-01; 1OT3OD025459-01; 1OT3OD025467-01, 1OT3OD025462-01), and from the Wellcome Trust (212930/Z/18/Z; 208381/A/17/Z). MD is supported by grants from NWO (400.17.605;628.011.011), NIH (3OT3TR002027-01S1; 1OT3OD025467-01; 1OT3OD025464-01), and ELIXIR, the research infrastructure for life-science data. MP was supported by the UPM Isaac Peral/Marie Curie cofund, and funding from the Dutch Techcenter for Life Sciences DP. LOBS and ES are supported by the Dutch Ministry of Education, Culture and Science (Ministerie van Onderwijs, Cultuur en Wetenschap), Netherlands Organisation for Scientific Research (Nederlandse Organisatie voor Wetenschappelijk Onderzoek), and the Dutch TechCenter for Life Sciences. We thank the NBDC/DBCLS BioHackathon series where many of these metrics were designed, and we especially wish to acknowledge the support and participation of the Dataverse team at IQSS, Harvard, and Todd Vision from Data Dryad.

